# Selection by differential survival among marine animals in the Phanerozoic

**DOI:** 10.1101/2023.09.21.558876

**Authors:** Erik Tamre, Christopher Parsons

## Abstract

The Gaia hypothesis posits that Earth’s biosphere functions as a single self-stabilizing system, but a key challenge is explaining how this could have arisen through Darwinian evolution. One theory is that of “selection by differential survival”, in which a clade’s extinction probability decreases with age as it accumulates adaptations resisting environmental disturbances. While this is hard to assess during early Earth history, we can assess whether this process operated among marine animal genera throughout the Phanerozoic. To that end, we analyzed time ranges of 36,117 extinct animal genera using fossil occurrence data from the Paleobiology Database in order to calculate marine metazoan extinction age selectivity, extinction rates, and speciation rates over the Phanerozoic. We identify four signatures of selection by differential survival: lower extinction rates among older lineages, heritability and taxonomically nested propagation of extinction resistance, reduced age selectivity during rare environmental perturbations, and differential extinction rather than speciation as the primary driver of the phenomenon. Evidence for this process at lower taxonomic levels also implies its possibility for life as a whole – indeed, the possibility of Gaia.

## Introduction

The Gaia hypothesis views the Earth’s biosphere, and indeed the entire Earth, as a self-stabilizing system promoting its own continuity [1]. Since its origin, a key challenge for this theory has been the difficulty of identifying a means by which such a system would be produced as a result of Darwinian evolution [2] [3] [4]. Recently, Doolittle [5] advanced a model of selection by differential survival (SDS), which provides a mechanism by which a Gaian biosphere could potentially develop and sustain itself. Under this evolutionary model, non-competing lineages occasionally develop adaptations that help resist environmental disturbances. Those that manage to persist for longer are thus likely to accumulate survival-enhancing adaptations – increasing age would lead to progressively lower extinction probability. Among lineages on the early Earth, the one leading to the last universal common ancestor (LUCA) may have won out by such a process, lending the Earth’s subsequent biosphere a degree of Gaia-type resilience [6] [7]. The potential for SDS to contribute to a Gaian biosphere has been appreciated [9], and models of varying complexity have been advanced to demonstrate how this mechanism can select for resilient biospheres [10] and develop stable biogeochemical cycles [11]. However, there is practically no record of the Earth’s earliest biosphere and there are no known extraterrestrial biospheres, so we cannot directly assess if SDS actually operated in an early planetary context.

Even so, is there evidence of such a process occurring during the more recent history of the Earth’s biosphere? If generally present across the entire history of life, it could still be detectable on the smaller taxonomic scales and shorter time scales that we can access, with clades smaller than all life under selection. Observing a history of SDS across many fossil lineages would lend potential to its operation in life as a whole. Using fossil occurrence data for marine metazoans from the Paleobiology Database (PBDB) [12], we test whether Phanerozoic patterns of extinction are consistent with a model of SDS. While Doolittle [5] was agnostic to how traits are propagated along lineages – considering them immortal “individuals” – propagation of traits across generations of an animal lineage proceeds through Darwinian inheritance. If inheritance is a substantial component of SDS in marine metazoans (potentially in addition to non-heritable traits), we would expect to see:

### Decreasing probability of extinction for a clade as it gets older

If, by persisting, clades evolve to become *more* persistent, we expect old clades to go extinct at lower rates than young clades. Such positive age selectivity of extinction has been previously observed [13] [14] [15] [16] [17], but no connection with SDS has been proposed.

### Heritability and taxonomically nested propagation of extinction resistance

We would also expect *subclades* of old clades to go extinct at lower rates than *subclades* of young clades. For example, a genus would benefit not only from its own age, but also the age of its order, since the accumulation of and selection for useful adaptations *sensu* Doolittle would happen along any branch in the tree, not only those defining certain taxonomic levels. Despite its non-Darwinian mechanism of selection, SDS in this context relies on heritable traits, which serve as “memory” of past events. Non-heritable traits such as geographic range and richness – though potentially useful for supporting a clade’s survival – should not confer any extinction resistance to constituent subclades.

### Reduced age selectivity during unprecedented environmental pressures

Age selectivity of extinction should diminish at time horizons when a novel selection pressure is imposed, as there would have been no past accumulation of adaptations tailored to that particular pressure. Major mass extinctions may be such horizons.

### Age selectivity “through survival alone”

Older lineages should persist because they experience a reduced rate of extinction, rather than increased rate of speciation.

Note that some of these features are in contradiction with the “law” of constant extinction and the Red Queen model of evolution that predicts it [18]. However, other researchers have offered different interpretations and methodological approaches [19] [20] [21] [22] [23] or suggested that this “law” does not apply on geological time scales [24]. Instead, a clade’s long-term fate may be determined more by interaction with the environment – such as proposed in the alternative Court Jester model [24] – and a level of environmental stability over time might allow for adaptations to be developed and selected that enhance the survivorship of older clades.

In an effort to find evidence for or against SDS among Phanerozoic metazoan lineages, we examine the time ranges of 36,117 marine metazoan genera to measure extinction rates, speciation rates, and age selectivity using fossil occurrence data from the PBDB. Note that SDS, given its non-Darwinian mechanism, could in principle also be observed in non-biological systems where lineages (companies, governments, or languages, perhaps) persist through time and interact with the environment – but we have chosen to focus on the biological system owing to its relevance to Gaian discourse and the ready availability of a thorough occurrence database. Our methods build in part on those of Finnegan et al. [13], but we significantly expand upon that study in order to discriminate between potential underlying evolutionary mechanisms. We identify all four signatures of SDS predicted above, providing the first empirical evidence that Doolittle’s theoretical framework describes a genuine evolutionary phenomenon in Earth history.

## Results

### Time-integrated extinction age selectivity

After downloading fossil occurrence data for 36,117 marine metazoan genera from the PBDB, we calculated age-dependent survival rates following [13], integrating over time and all genera (Fig. 1a). As expected, young genera exhibit much higher extinction rates than old genera. Slightly less than 50% of the youngest genera appear in the next 11-Myr time bin, while 80% of genera survive the transition from the fourth time bin into the fifth (at 44 Myr).

**Fig. 1:**
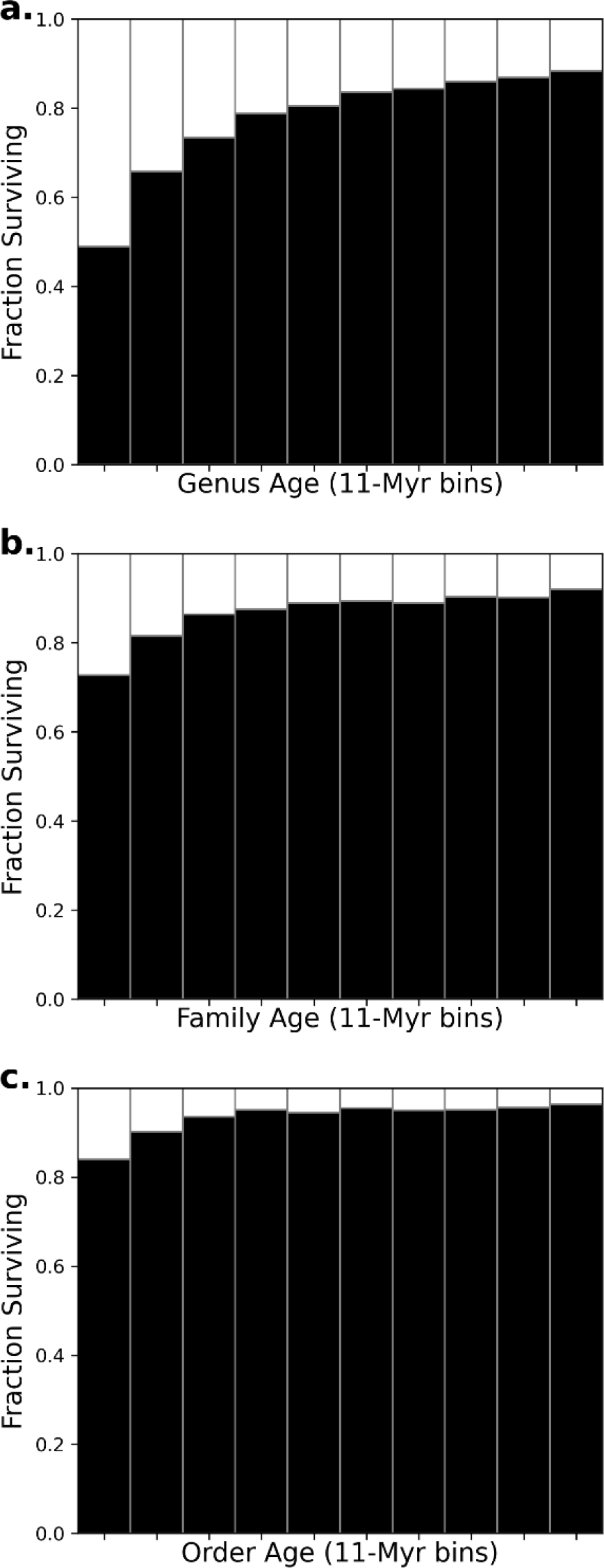
Age selectivity of extinction at different taxonomic levels in marine metazoans. Time-integrated survival probability as a function of clade age compiled for genera (a), families (b), and orders (c). Clade age is measured in 11-Myr time bins, and survival rate is expressed as the fraction of taxa that reoccur in the subsequent time bin.

Increasing age grants progressively diminishing increases in survival probability, and after 110 Myr, the chance of survival into the next bin saturates at about 90%. We further tested the same effect at the level of families and classes (Fig. 1b-c). While families experience lower extinction rates than genera, and orders lower still, both distributions display clear, though diminished, extinction age selectivity.

Using logistic regression (Online Methods), we inferred an 11% multiplicative increase in generic survival probability per 11 Myr (6.6% for families and 5.8% for orders). However, the aforementioned saturation effect complicates the fit, as logistic regression assumes a constant benefit provided per unit increase in age, unlike these distributions in which young taxa receive greater benefit from aging than already-old taxa. In effect, these percentages are a compromise between the unsaturated and saturated fitness effects (Extended Data Fig. 1).

To exclude misspelled occurrences from the dataset, we removed genera that could not be assigned to a known taxonomic order, as misspellings would appear as short-lived genera represented by a single occurrence, artificially decreasing the survival rates of one-bin-old taxa. Even after this filter, the short-lived taxa in our dataset presumably still include some misidentified or improperly logged entries. However, any genuine short-lived genus would also be expected to have few, if any, occurrences, either due to local ecological distribution, preservational bias, or indeed their short time on Earth. Filtering our dataset to exclusively include genera supported by multiple temporally-unique occurrences only biases one-bin-old survival rates in the opposite direction: survival for one-bin-old genera increases by 15% if only genera with multiple occurrences are included, and by another 15% if those occurrences must be dated to different times. This operation brings one-bin-old taxa to an unreasonably high survival rate (>80%), which implies that most of the discarded data were valid (Extended Data Fig. 2). However, we observe that any misclassified occurrences remaining in the dataset will artificially increase the strength of the inferred age selectivity.

Additionally, to ensure that extinction resistance in older genera – a phenomenon demonstrated in Metazoa as a whole – is not being driven by a clade-specific bias, we subdivided the dataset into various major animal phyla, and recalculated age selectivity. Together, Mollusca, Arthropoda, Chordata, and Brachiopoda comprise 81% of all marine animal genera in the PBDB (32%, 20%, 19%, and 9%, respectively; Extended Data Fig. 3). In each phylum, older genera reliably go extinct at lower rates than younger genera, recapitulating the aggregate pattern of age selectivity and establishing the persistence effect as a phenomenon operating across a wide range of metazoan evolutionary trajectories (Fig. 2).

**Fig. 2:**
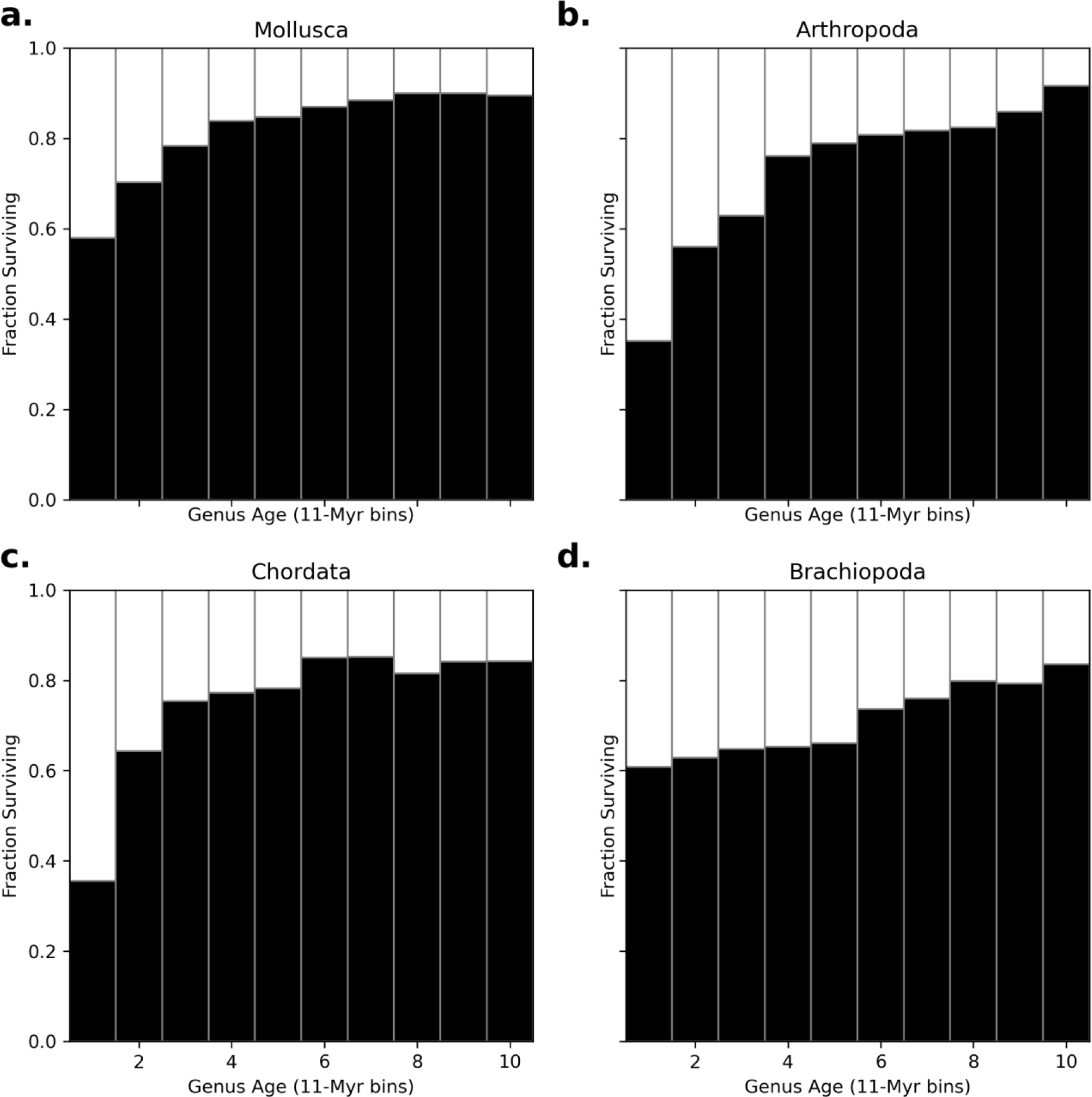
Age selectivity of generic extinction in major metazoan phyla. Time-integrated age selectivity curves for the four best-represented metazoan phyla in the dataset: Mollusca (a; 31% of genera), Arthropoda (b; 20% of genera), Chordata (c; 19% of genera), and Brachiopoda (d; 9% of genera).

### Time-dependent data for all Metazoa

Beyond a detectable persistence effect within aggregated datasets, we also observe that the strength of age selectivity has varied over geologic time. The simplest proof of this variance is that mean genus age has not monotonically increased over the course of metazoan evolution (Extended Data Fig. 4). We divided the period between 550 Ma and the present into one hundred 5.5 Myr bins and measured the strength of persistence across each bin boundary by fitting logistic regressions to the observed extinction rates for genera of varying age (Fig. 3). Following previous work [13], our metric for the strength of age selectivity is the natural log of the rate parameter of the fitted model; positive values indicate positive (old-favoring) age selectivity and negative values indicate negative (young-favoring) age selectivity. While the strength of age selectivity still fluctuates over geologic time, it is almost always positive (i.e. old genera usually experience lower extinction rates than young genera). However, at certain time points, notably those corresponding to the four most recent canonical mass extinction events, log-odds approach zero, implying a temporarily age-independent extinction pattern. Importantly, the log-odds global minimum (value) occurs at precisely the End Permian boundary. Additionally, following the pronounced local minima at the End Permian, End Triassic, and End Cretaceous, we observe a rebound effect, in which the reimposed age selectivity is stronger than it was prior to each extinction. Extinction age selectivity and speciation rate are weakly negatively correlated, and this correlation is absent under low extinction rates (Extended Data Fig. 5). Despite the low correlation, there are no instances of a high extinction rate coinciding with strong age selectivity; the data define a clear linear boundary in which age selectivity is constrained.

**Fig. 3:**
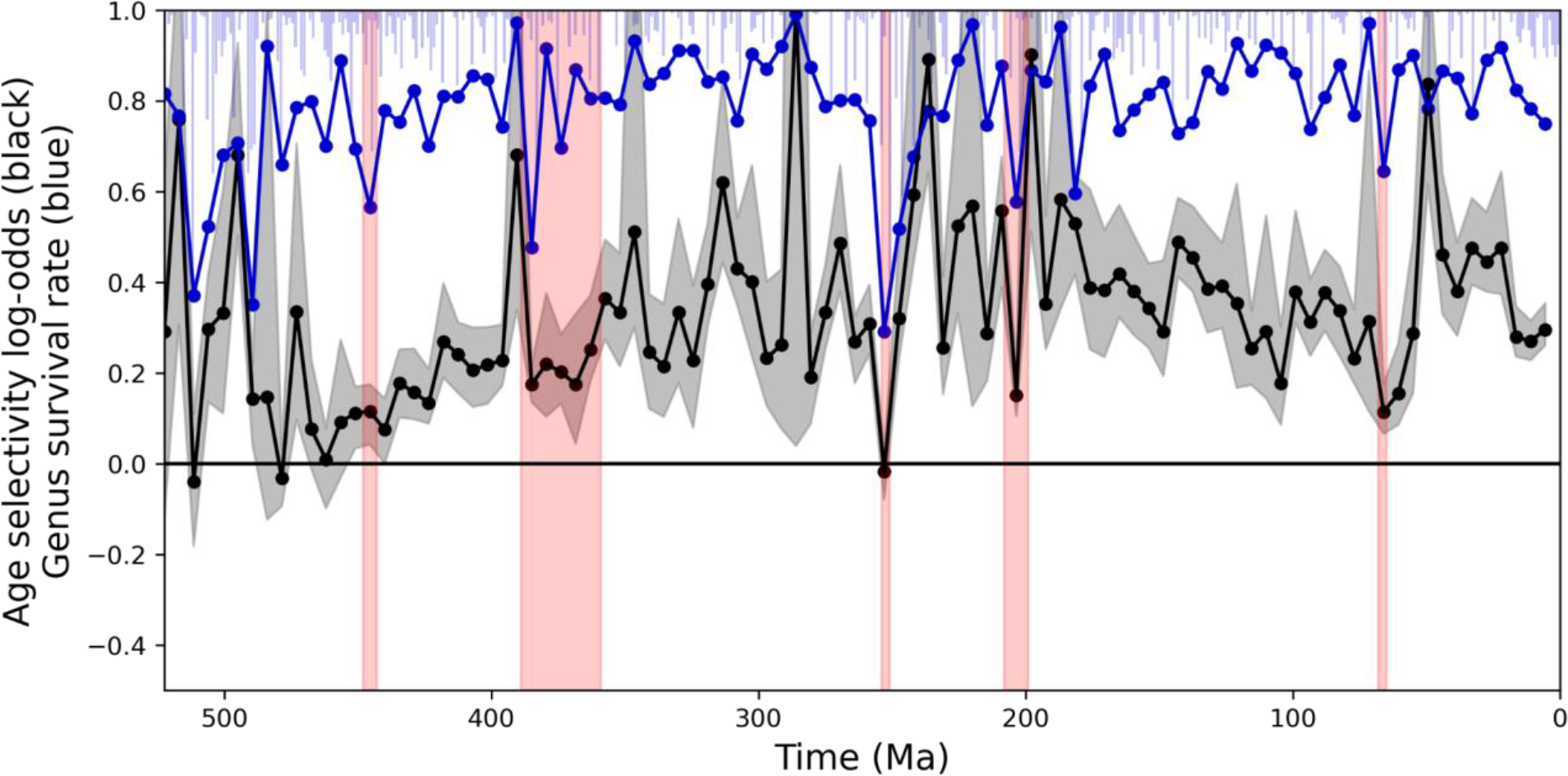
Survival rate and age selectivity of extinction and for marine metazoan genera over the Phanerozoic. Survival rate is plotted in blue and age selectivity in black. Each dot represents a 5.5 Myr time bin boundary. Age selectivity log-odds values express the strength and direction of the age selectivity of extinction, with positive values implying higher survival rates for older genera, negative implying the inverse, and near-zero values implying age-independence. Uncertainty was calculated from 100 bootstrap replicates of the population across each bin boundary, and is denoted by the gray shaded region. The light blue bars depict extinction rate with a higher time resolution, to show any patterns obscured by coarse binning. The 5 canonical mass extinctions (Late Ordovician, Late Devonian, End-Permian, End-Triassic, and End-Cretaceous) are shown in red.

### Heritability of extinction resistance

In an effort to disentangle the obvious extinction-buffering afforded by a clade’s richness and range from the subtle extinction resistance that may be conferred through heritable traits, we divided our observations of generic extinction and speciation based on the age or richness of the containing order at the time of observation. The key idea of this methodology is that, while an individual genus is blind to the richness of its containing order, it is not necessarily blind to its order’s antiquity. If persistence-enhancing traits do exist, and are sufficiently retained across genera within an order, genera in old orders are expected to experience less extinction than genera in young orders, even if the genera themselves are of equivalent ages. Genera in rich orders, however, are only expected to have enhanced survival rates to the limited extent that a clade’s richness is correlated with its age [25] (e.g., Krug et al., 2008), as richness provides no fitness benefit to a clade’s individual members. While noting that generic richness and clade range can be associated as well, we do not attempt to directly estimate range, as range data in PBDB is sparse and especially prone to taphonomic bias.

Strikingly, order age cleanly divides trends of generic extinction, speciation, and even age selectivity of extinction, with genera in old orders experiencing much lower extinction rates, speciation rates, and extinction age selectivity (Fig. 4). Comparatively, richness inconsistently divides these metrics, and only ever weakly. Nevertheless, rich orders and old orders *themselves* experience substantially lower rates of extinction than young orders and poor orders. Taken as a whole, we observe that while both a clade’s age and generic richness increase its chance of survival, only the clade’s age improves the survival chances of its constituent subclades. This discrepancy suggests that the fitness benefit provided by age is heritable, unlike that provided by richness. The same trend is observed for genera divided according to family and class (Extended Data Figs. 6 and 7) instead of order.

**Fig. 4:**
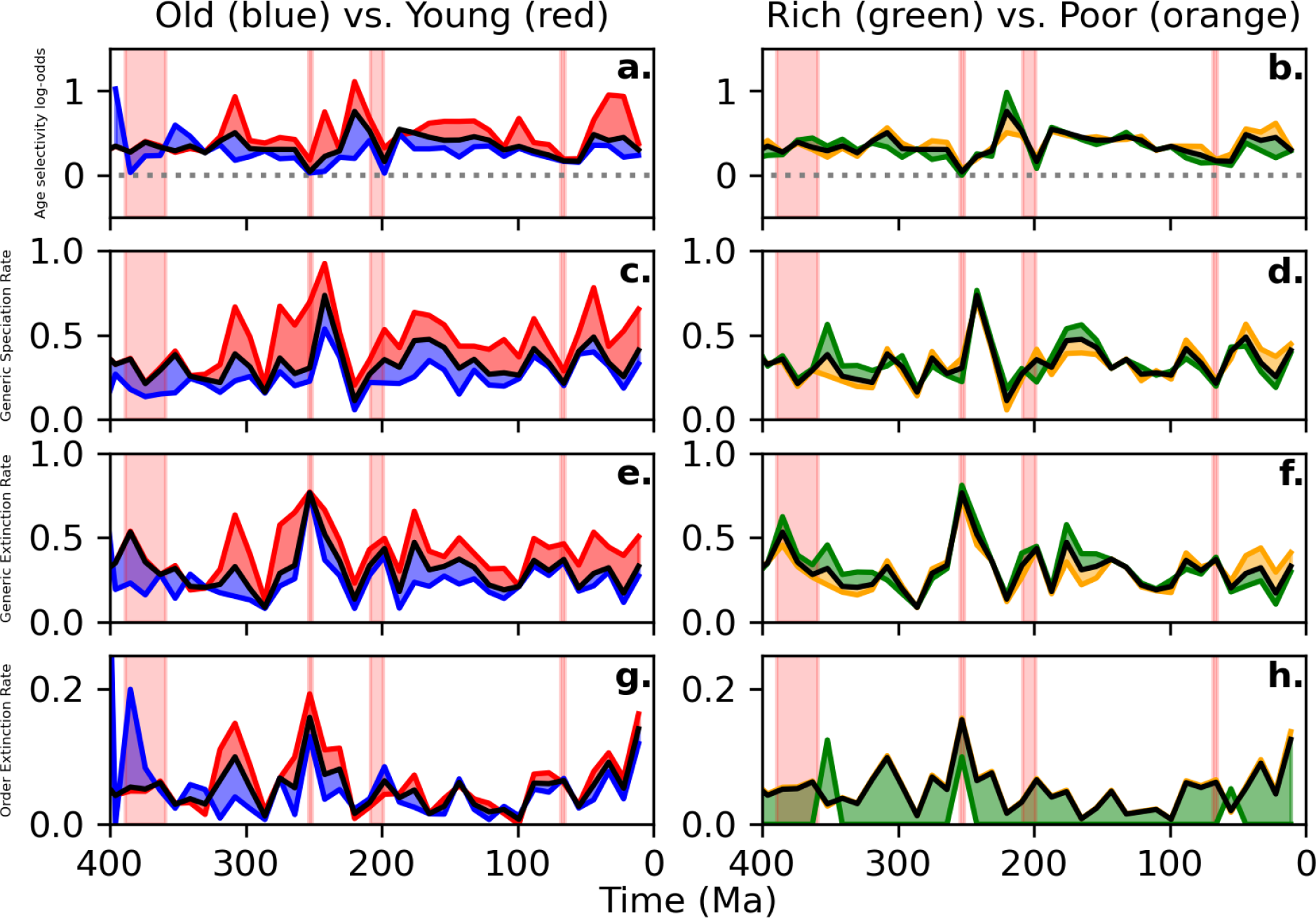
Extinction dynamics divided by taxonomic order’s age and generic richness. Four evolutionary metrics plotted over the duration of the Phanerozoic: Age selectivity of extinction (a-b), generic speciation rate (c-d), generic extinction rate (e-f), and order extinction rate (g-h). At each bin boundary, genera (or orders themselves in g-h) were divided into two groups – by the *age* of their order in the left column, and by the *generic richness* of their order in the right column. Orders are defined as “old” if they are older than 11 time bins (143 Myr), and “rich” if they contain more than 40 genera. The color scheme is “old”: blue, “young”: red, “rich”: green, and “poor”: orange. The black line in each panel is the value of the metric for the undivided dataset; note that the metrics differ more clearly and consistently when split by age rather than generic richness. Data are reported from 400 Ma onward, as most metazoan orders appear in the fossil record after 550 Ma, and therefore prior to 400 Ma the existence of an “old” order is impossible. The Late Devonian, End-Permian, End-Triassic, and End-Cretaceous extinction events are shown in red.

## Discussion

In his initial outline of SDS, Doolittle [5] presents the process in its distilled form: even in the complete absence of competition, evolving lineages can accumulate useful traits that incidentally confer resistance, physiological or otherwise, to environmental fluctuations. Many such traits are likely to be subtle, composite, and difficult to observe in the fossil record.

Persistence-conferring traits need not be strictly Darwinian either: non-heritable traits such as species richness, ecospace coverage, and geographic range would undeniably shield a large clade from extinction. Therefore, our observation that rich clades rarely go extinct is to be expected; a rich clade must succumb in a series of extinction events, by the end of which it will, by definition, no longer be a rich clade. Importantly, we show that the genera *within* rich clades do not experience significantly different rates of extinction, speciation, or age selectivity than genera in poor clades (see Fig. 4d), implying, as expected, that generic richness is beneficial to the collective, but not to constituent subclades.

Instead, clades *and their individual subclades* benefit from increased age of the clade (see Fig. 4c). If age and richness are sufficiently correlated (i.e., if clades tend to increase in richness as they age), the observation that old clades experience diminished extinction rates could simply be attributed to their greater richness. In that case – if richness determines survivability – survivability would not be heritable. However, if Darwinian traits also contribute, survivability should be heritable to some degree: if a family, order, or class is sufficiently old, the genera within should be more extinction-resistant than genera in a young family, order, or class. In other words, a genus might inherit some persistence-enhancing traits as an ancestral character. Indeed, it is one of our key observations that genera in old families, orders, and classes suffer markedly lower extinction rates than those in young families, orders, and classes – while the generic richness of the family, order, or class is not associated with a substantial change in generic extinction rate. This suggests that some persistence-inducing traits are Darwinian in nature, accumulating as a lineage ages and experiences a diversity of environmental conditions and climate events.

We also observe a reduction in age selectivity at the genus level during large-scale extinction events (Fig. 3): the advantage of old genera over younger ones vanishes or diminishes at these horizons. This is what a model assuming that adaptations accumulate over time in response to consistent environmental pressures would predict, as few lineages would have been tested for the near-unprecedented conditions during major extinction events. However, the generic extinction rate remains higher in young orders than in old orders throughout the record, even during mass extinctions (excepting the P-T at the order level; Extended Data Fig. 7e suggests that a gap in extinction rate might persist even there at the class level). Perhaps orders and classes, unlike genera, live long enough to have experienced previous mass extinction-like horizons and their synapomorphies have thus been influenced by these selection pressures, conferring their members a degree of persistence during subsequent events. One might say that the Court Jester (Benton, 2009), though unpredictable, remains partial to certain jokes. For example, an emplacement of a large igneous province has occurred many times over the Phanerozoic with somewhat predictable ecological consequences [26], and a suggestion of high-level clades having developed some long-term resistance to these repeated hazards is perhaps as close to Doolittle’s planetary vision as we can empirically demonstrate.

Do extinction resistance traits accumulate at any particular taxonomic level? We would argue that they are accumulating along all branches of the metazoan tree simultaneously, and that such traits can be acquired by any evolving lineage. The fact that family, order, and class age all increase genus survivability (Fig. 4c; Extended Data Figs. 7 and 9) means that not only is extinction resistance heritable, but also that these heritable traits can appear along branches of any depth in the tree. While we did not study the effect of genus age on species survivability, our results suggest that even a genus can gain additional traits to resist environmental challenges over the course of its duration. If that genus continues to thrive and diversify, it may grow in diversity to the point that it would be better classified as a family, and so on. This creates a recursive effect, in which a clade’s retained characters from all ancestors supplement those generated as the clade persists and evolves. Yet not all ancestral traits will be relevant to descendant lineages, especially if the descendant’s lifestyle or the Earth system itself changes substantially. While our results suggest that extinction resistance for a single genus tends to saturate in <110 Ma; Fig. 1a), we still observe that the overall age selectivity of extinction has not grown discernibly weaker over a period five times the apparent saturation time (Fig. 3). This may be because lineages need to adapt to a moving target, which requires not only the retention of useful ancestral traits, but also the acquisition of new traits in response to competition and the changing environment.

Our results demonstrate that a process enhancing the survival probability of older clades is operating across a variety of taxonomic scales, across different groups among the Metazoa, and throughout Phanerozoic time. The robustness of the observed effect with respect to slicing the dataset in these ways helps exclude a well-known statistical artifact producing illusory age selectivity owing to mixture of differing extinction rates across taxonomic groups and time (see [13] for a detailed review). It also reduces the likelihood that our inferences are dominated by inaccuracies in interpreting the capricious paleontological record – characterized by taphonomic bias, dating uncertainty, and imperfect correspondence between assigned taxon names and monophyletic groups or continuous lineages (see Online Methods for treatment of raw data).

Through differential survival, animal clades can thus evolve to persist in the face of environmental fluctuation. If, as Doolittle [5] suggests, the cumulative effect of SDS on a planetary scale may have been a self-stabilizing *biosphere*, it is also natural to ask if animals themselves may have any substantive part to play in it. In many ways, animal activity does contribute to homeostasis of the environment [27]: marine calcifiers make their skeletons out of carbonate which partakes in the buffer system enhancing the ocean’s carbon dioxide uptake [28], bioturbation stabilizes the carbon cycle by mixing the sea floor [29], and heterotrophy by zooplankton acts as a control on algal blooms [30]. This is not to say that animals are responsible for any possible Gaian properties of the system (which would have been established billions of years before their emergence), but that, like all lineages, even animals can contribute to overall stability.

On a process level, the scale-independent nature of the observed persistence effect suggests a possible extrapolation beyond the range of taxonomic scales here considered to life as a whole. Furthermore, life at its early stage might have been equivalent to an order or class in terms of the evolutionary distances between its members anyway, removing any need for such propagation. For a possible mechanism for Gaian life to become established on the early Earth, it is also noteworthy that we do not identify richness or range as chief variables conferring persistence – it is genuinely associated with a taxon’s age and seems to involve heritable traits. That is what would be necessary for life to persist on a planetary scale: richness and range would not have helped early life, but a planetary-level threat would rather have required stabilizing adaptations. For example, a proposed early “Gaian bottleneck” owing to the accumulation of gases leading to a runaway greenhouse effect [31] would have required an adaptation allowing for the biological regulation of atmospheric composition, so as to prevent planetary extinction. Our detailed observations of survivorship outcomes in the Phanerozoic suggest an underlying continuous process that is consistent with Doolittle’s view of what might have been at work on the early Earth. Whether it contributed to any emergence of a Gaian biosphere and its continued stability or not – an admittedly different planetary context – this evolutionary dynamic has been operating on smaller taxonomic scales through the Phanerozoic. Even if restricted to the directly observed scope, this has major implications for long-term survival of complex life – which is not a given, considering the history of mass extinctions in the Phanerozoic.

## Online Methods

### Extraction of data from PBDB

We downloaded occurrence data from the Paleobiology Database (Alroy et al., 2001), using the search terms *base_name*=“Metazoa”, *envtype*=“marine”, and *show*=“classext,acconly”. With custom Python scripts, we parsed the results, first calculating the midpoint of each occurrence’s possible age range and then binning the occurrences into the appropriate genera. Any genus that could not be mapped to a known order was excluded from the dataset, as such instances likely represent misspellings or misclassifications of accepted genera, making the record appear overly age-selective. Conversely, any genera that appeared to persist in the fossil record for longer than 250 Myr were also removed, as they likely represent wastebasket taxa, rather than true clades, and would therefore also unduly strengthen the signal of persistence.

By dividing the period from 550 to 0 Ma (the interval containing the vast majority of diagnostic metazoan fossils) into one hundred 5.5 Myr time bins, we compiled all occurrence data into a matrix, where each entry represents the age (in discrete time bins, starting at 1) of a particular genus at a particular point in time, or 0 if the genus has gone extinct or has yet to appear. A genus is assumed to have existed for the entire duration between its first and last occurrence, even if it does not appear in all intervening time bins. To generate bulk persistence results, we repeated this methodology using 11 Myr time bins and pooled the results. Bulk persistence results were also calculated for various Metazoan subclades by varying the *base_name* search term.

Duration matrices could then be converted to histories of extinction and speciation with a value reported for each time bin. To do so, any appearances of age = 1 taxa were treated as speciation events and any instances of a taxon transitioning from age > 0 to age = 0 across a bin boundary were logged as extinction events. Counts were normalized to rates by dividing by the total number of extant taxa at the time of the observation.

Further, we logged to which family, order, and class (hereafter referred to as superclades) each genus belonged and used that data to calculate generic richness of each genus’s superclades in each of our time bins, resulting in paired duration and richness matrices for each taxonomic resolution. No richness matrix was inferred at the genus level, as the generic richness of any genus is by definition equal to 1.

### Splitting generic observations by superclade age and richness

By applying the above procedure to derive duration and richness matrices for families, orders, and classes, we separated individual observations of generic extinction and speciation based on the age or generic richness of its containing superclade *at the time of the observation*. By tallying so, a genus can be counted as a genus in a young/poor superclade in an early time bin and later be counted as a genus in an old/rich superclade. Thresholds to classify genera into one category or another were tuned manually to approximately evenly split the total number of observations and are as follows: “young” families, orders, classes ≤ 55 Ma, 143 Ma, and 275 Ma, respectively; “poor” families, orders, classes ≤ 5 genera, 40 genera, and 200 genera, respectively (Extended Data Figs. 8 and 9).

### Calculation of logistic regressions

Following [13], at each boundary between two time bins, we observed which taxa survived into the next time bin and which apparently died out. As this is a problem of using a discrete variable (age) to estimate the probability of a binary outcome (survival), we optimized logistic regressions (Equation 1) to model the effect of genus age on survival probability, *p(x)*.

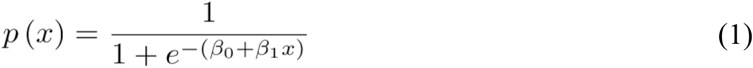

The term *β1* is known as the odds ratio or rate parameter, and in this case conveys the expected change in survival probability precipitated by a change in age. The natural log of *β1* yields the age selectivity log-odds, which we refer to as the age selectivity parameter. Regressions were conducted using the LogisticRegression module of scikit-learn [32] with default parameters. Bootstrap confidence intervals were calculated at each time bin by resampling the observations of extinction/survival 100 times and recalculating the regression parameters. As in [13], when modeling age selectivity, we only measured taxa less than 10 time bins old at each bin boundary, thereby truncating the occurrence interval of any taxon that exists for more than ten time bins to those first ten time bins. Therefore, by not counting the later occurrences of long-lived genera, we further mitigated potential bias, without removing such genera from the dataset altogether.

## Code availability

Code for the analyses performed in this study is available at https://github.com/DifferentialSurvival/SDS.

## Extended Data Figures

**Extended Data Fig. 1:**
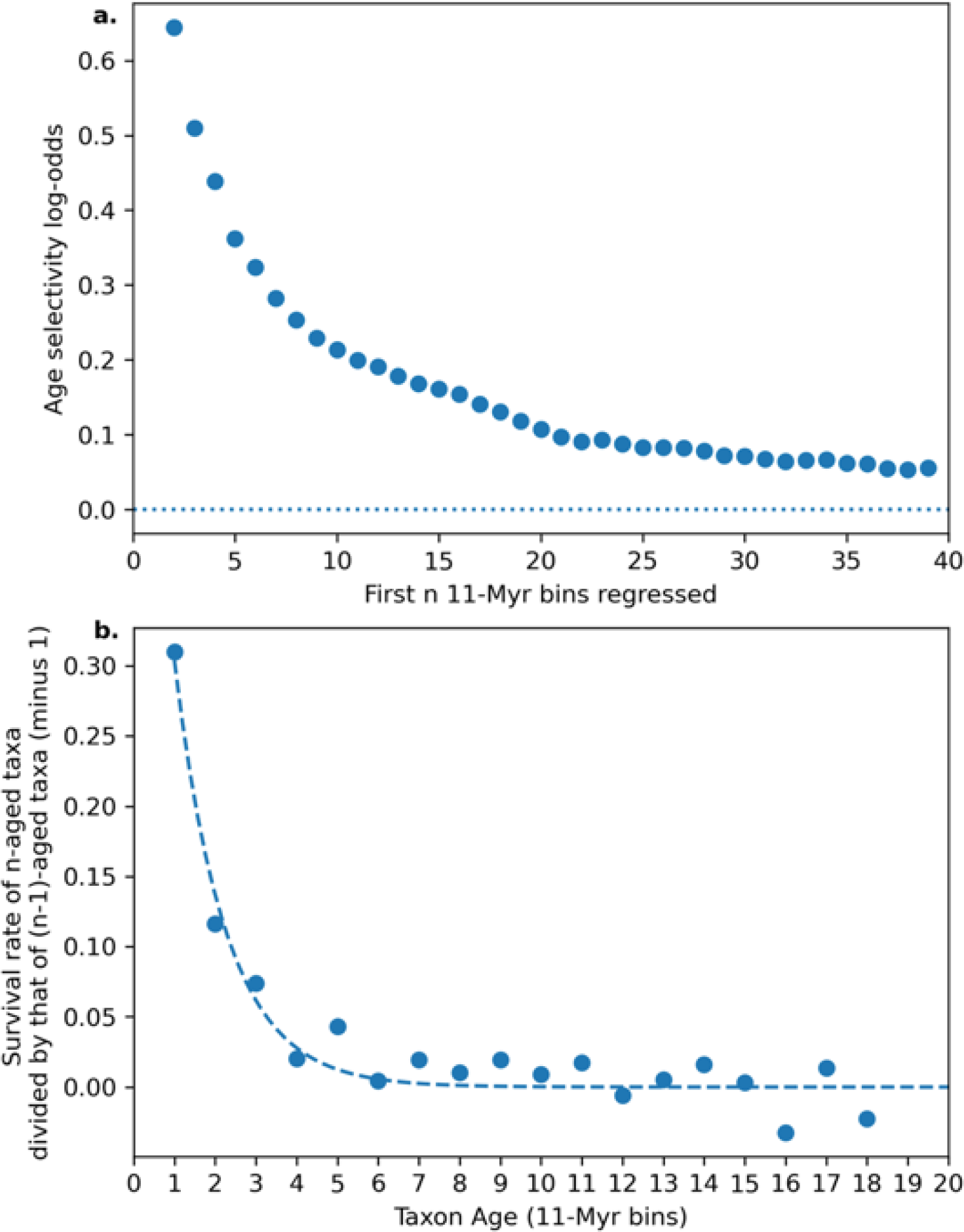
Decay of extinction risk reduction with increasing age for metazoan genera. In panel a, we fit a logistic regression on the data presented in Fig. 1, but with increasing numbers of age bins included. The absolute magnitude of age selectivity log-odds is therefore to some degree arbitrary, as it depends on how many age bins are considered. Similarly, panel b shows the “diminishing returns” of increased age. For example, aging from a 1-bin-old to a 2-bin-old taxon apparently provides a much larger survival benefit than aging from a 5-bin-old to a 6-bin-old taxon.

**Extended Data Fig. 2:**
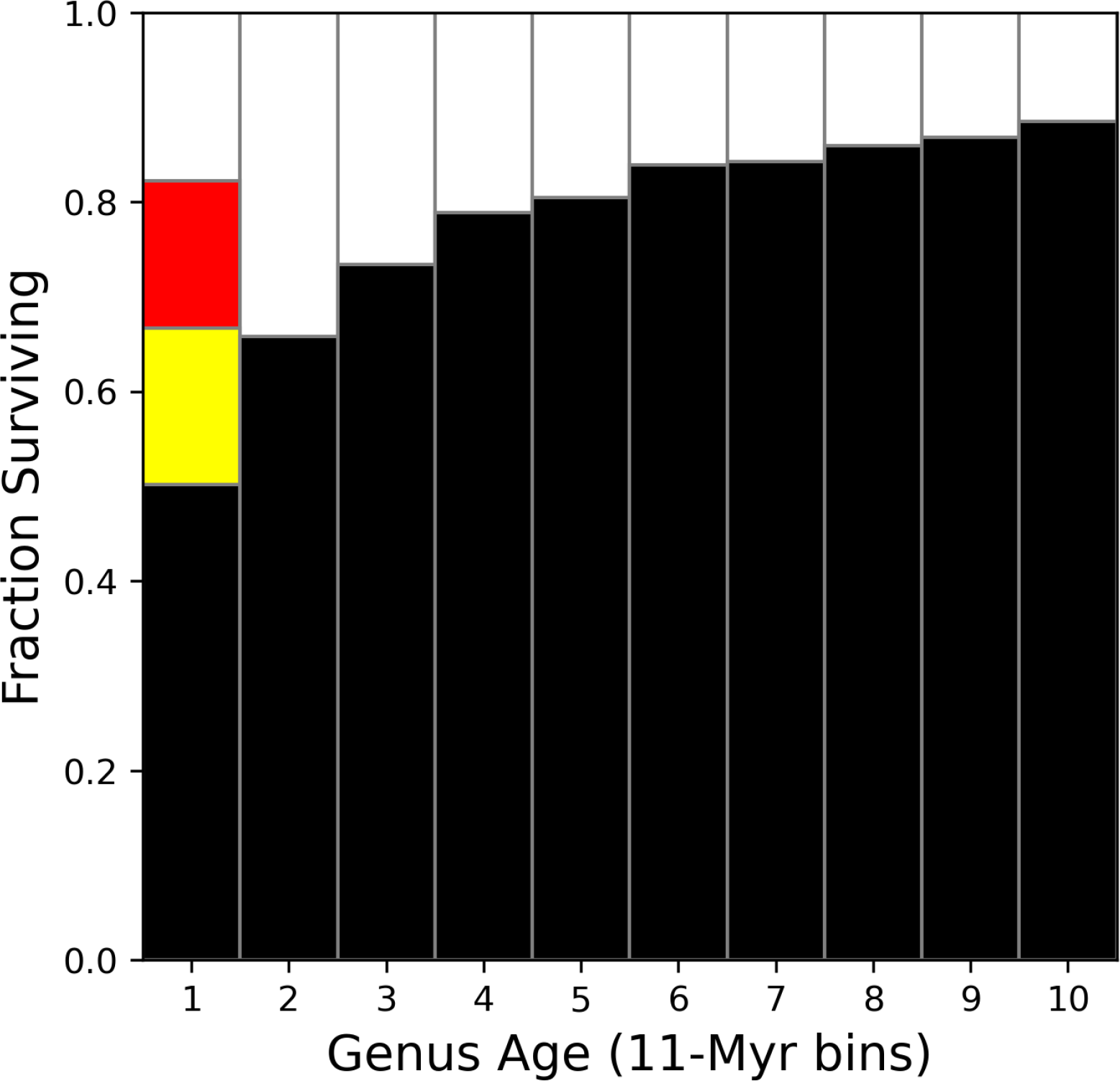
The aggregate effect on one-bin-old generic survival rates of only including genera with multiple occurrences (yellow) and of only including genera with multiple *temporally unique* occurrences (red).

**Extended Data Fig. 3:**
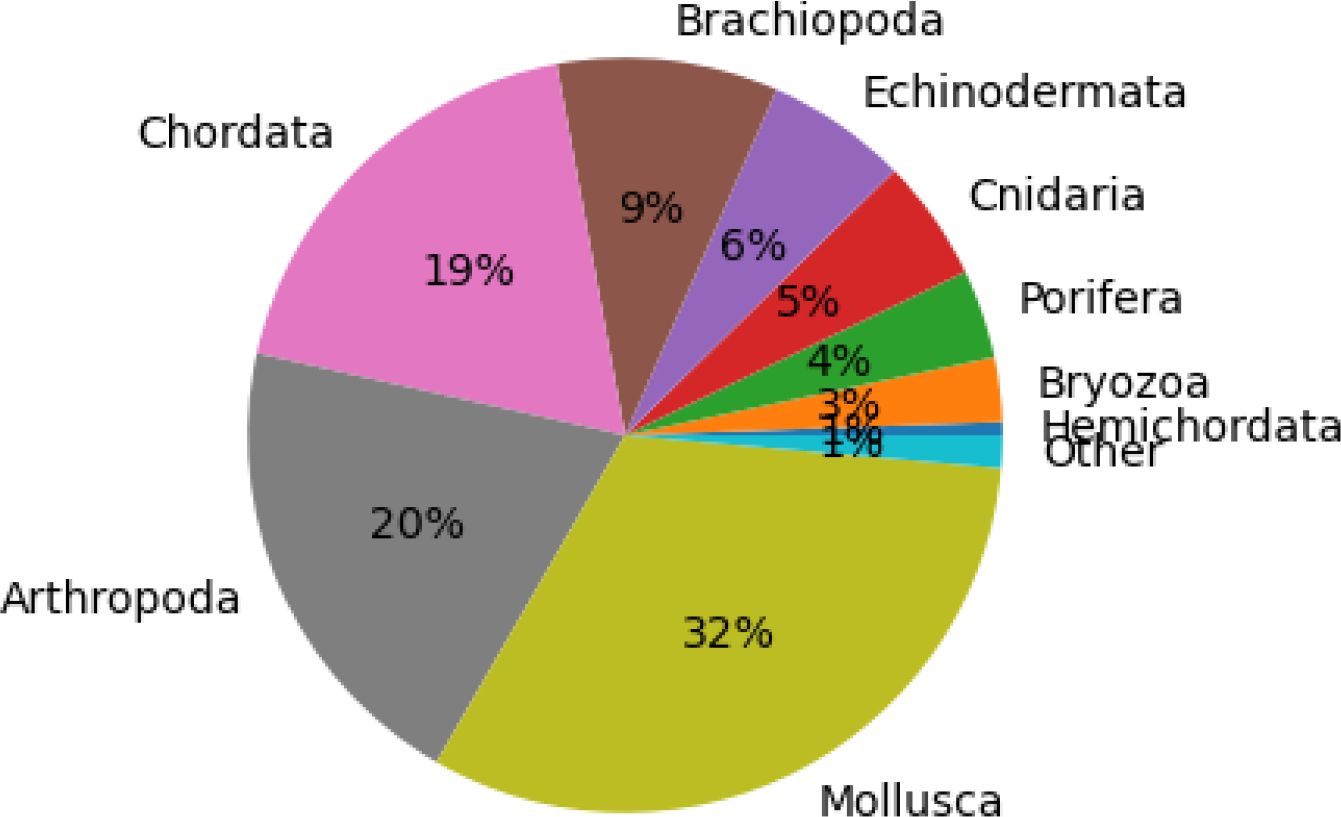
Phyletic composition of our dataset of 36,117 marine metazoan genera.

**Extended Data Fig. 4:**
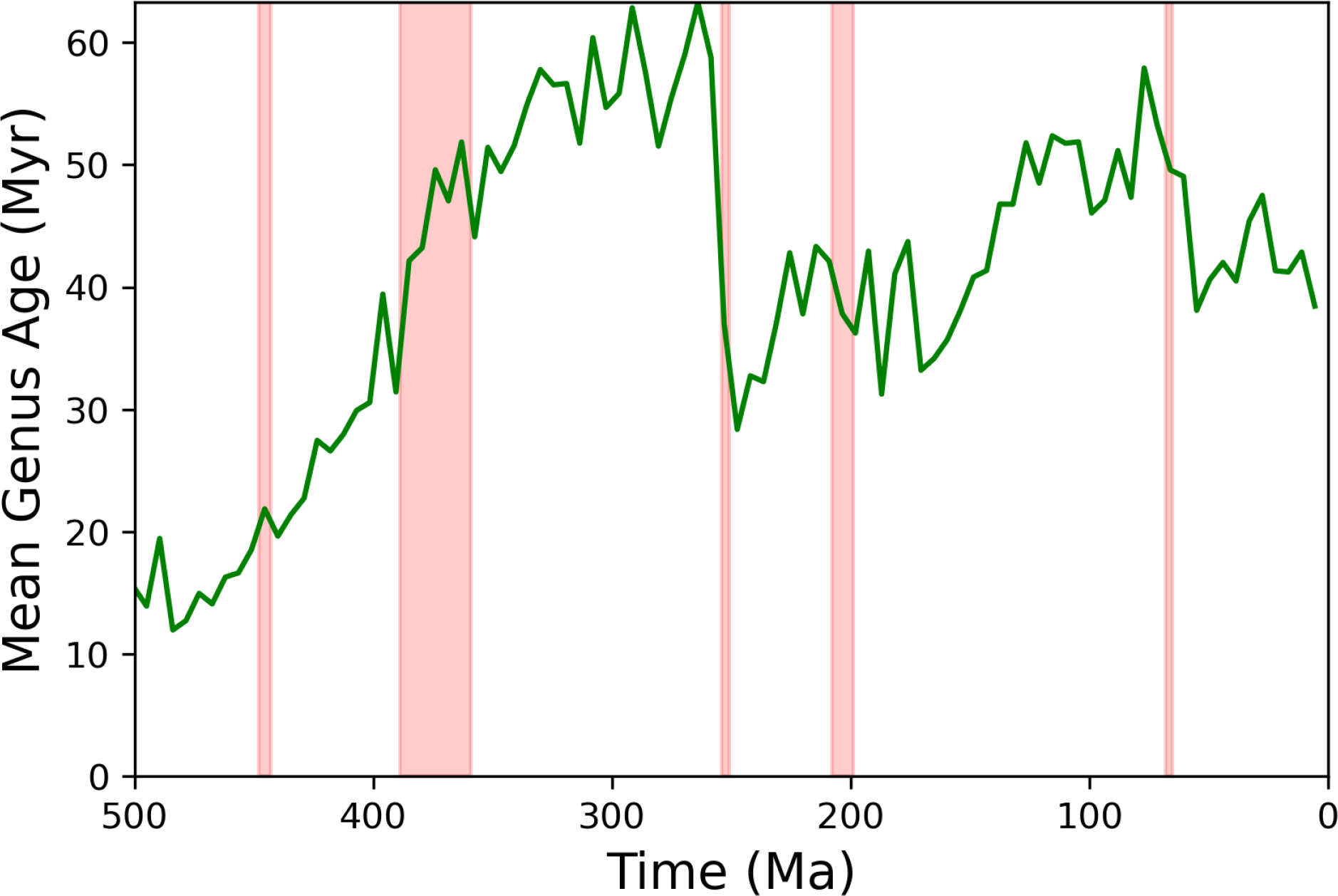
Mean marine metazoan genus age throughout Earth history. The 5 canonical mass extinctions (Late Ordovician, Late Devonian, End-Permian, End-Triassic, and End-Cretaceous) are shown in red.

**Extended Data Fig. 5:**
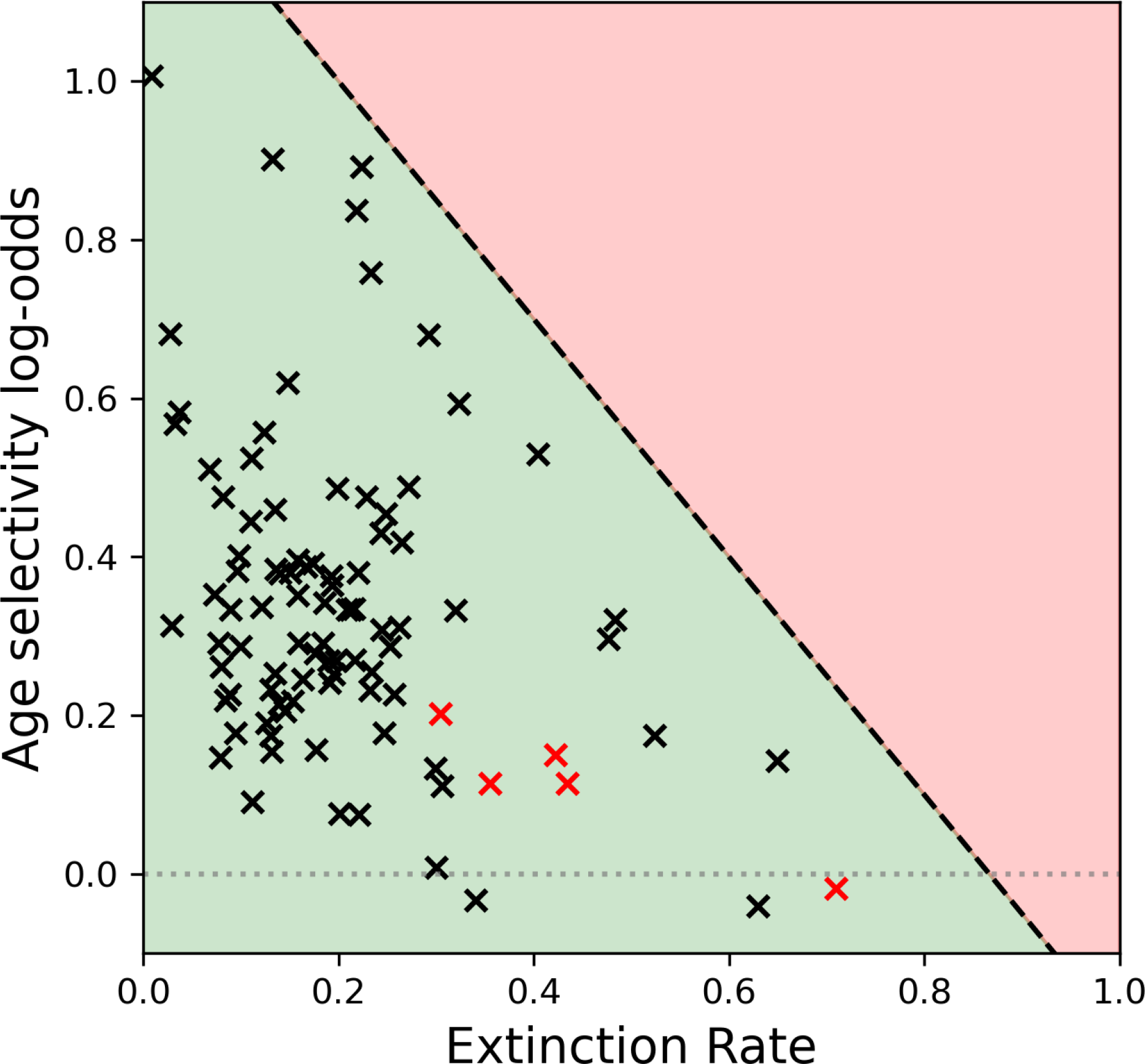
Extinction age selectivity log-odds plotted against extinction rates. Each point denotes a time bin boundary. The green shaded area roughly represents the observed range of extinction rate and age selectivity values, whereas the red area represents the apparent domain of incompatibility. For reference, bin boundaries corresponding to the five canonical major mass extinctions are shown in red. Note that they do not stand out extremely in this context, as a mass extinction is usually defined by a sizable net loss of diversity, which requires not only a high extinction rate, but also a low speciation rate.

**Extended Data Fig. 6:**
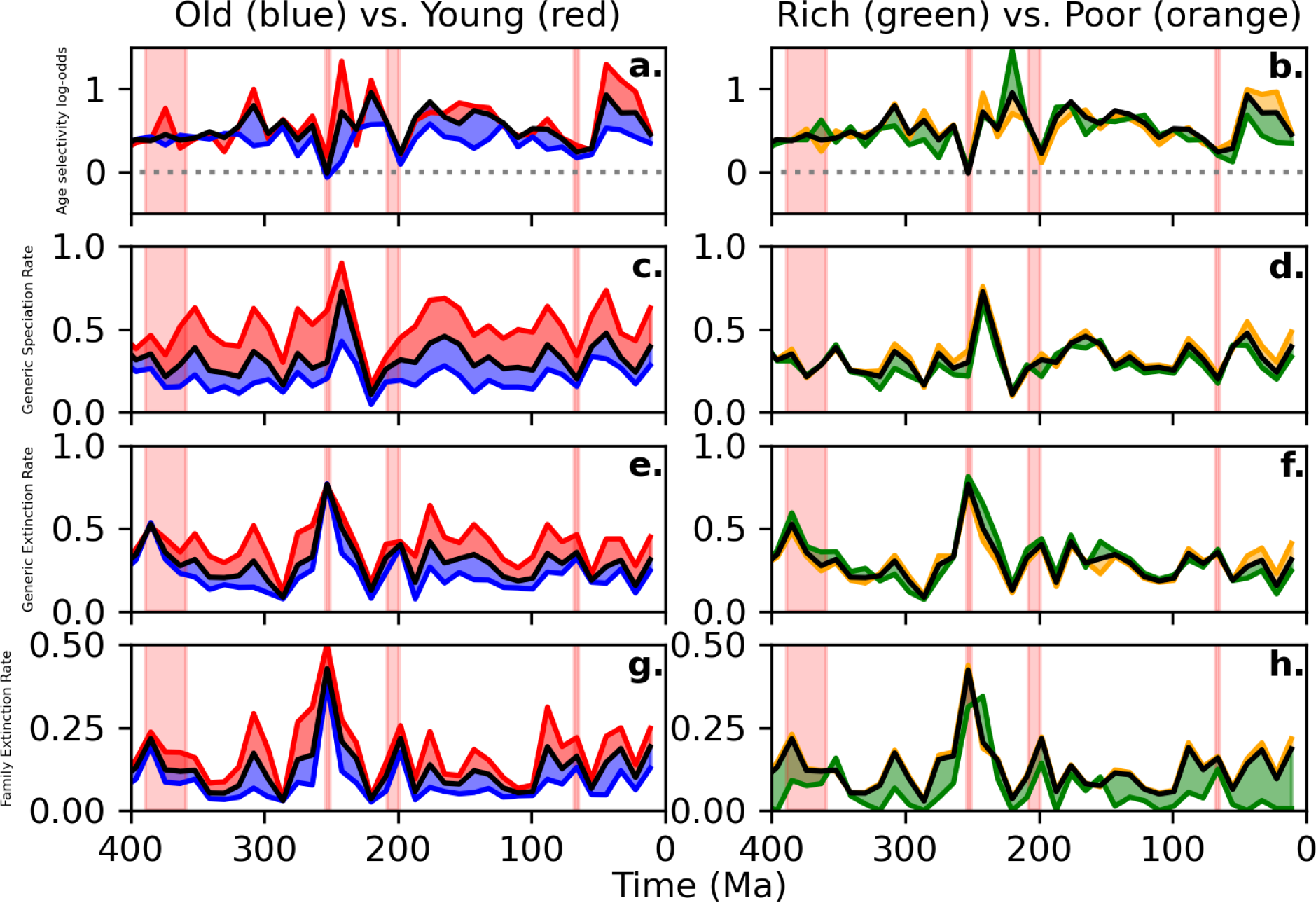
Extinction dynamics divided by taxonomic *family’s* age and generic richness. See Fig. 4 for a detailed explanation. Families are defined as “old” if they are older than 5 time bins (55 Myr), and “rich” if they contain more than 5 genera. While this figure highlights the same phenomena as in Fig. 4, inference of age selectivity for “young” families (panel a) proved complicated, as regressions were in general fitted using the first 10 age bins at each time point, but the young family cutoff should be substantially lower than 10 age bins to be informative. Therefore, the values reported in panel a were calculated using only the first 5 age bins, both in “young” and “old” families.

**Extended Data Fig. 7:**
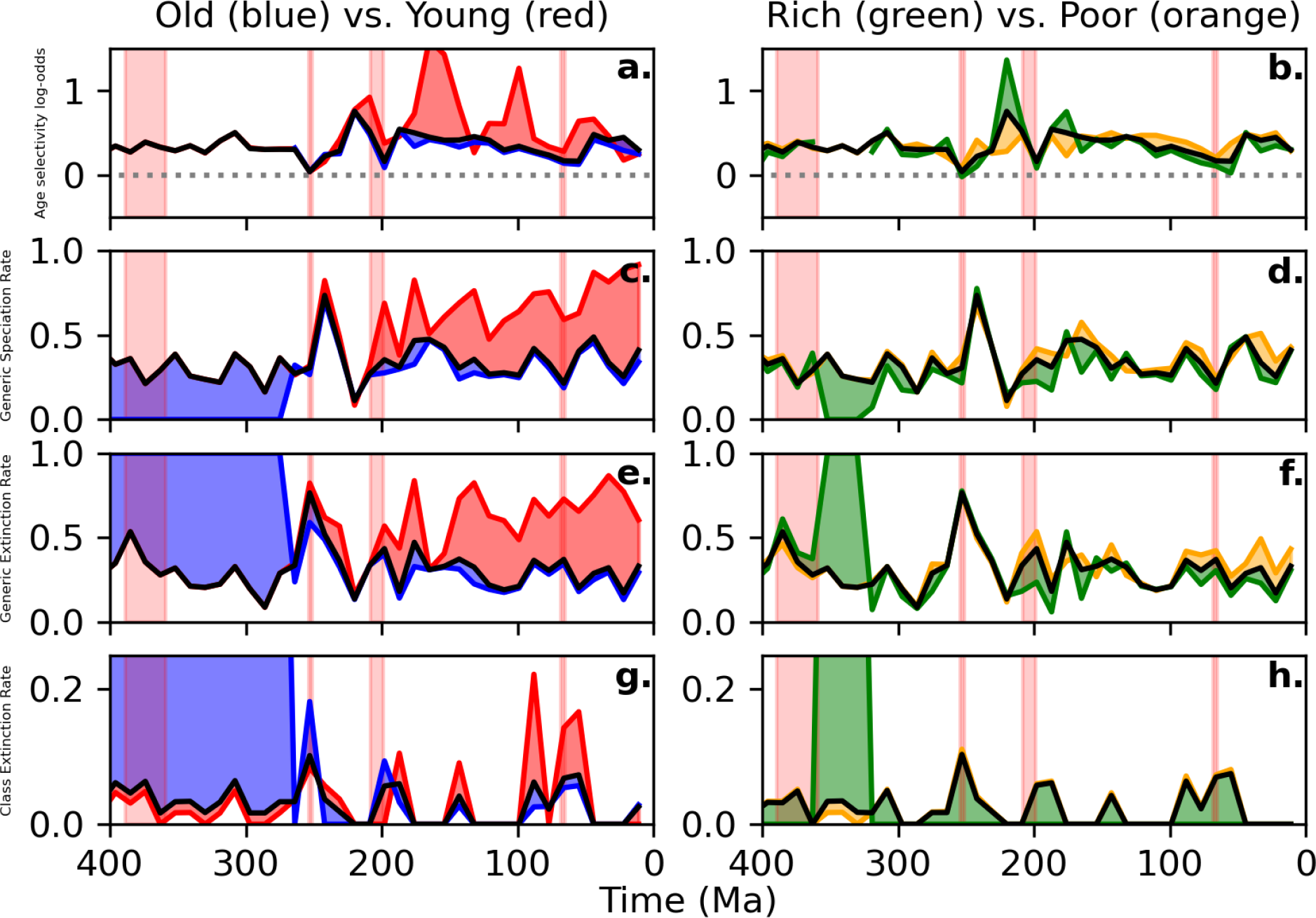
Extinction dynamics divided by taxonomic *class’s* age and generic richness. See Fig. 4 for a detailed explanation. Classes are defined as “old” if they are older than 25 time bins (275 Myr), and “rich” if they contain more than 200 genera. While this figure does to some degree recapitulate the effect we demonstrated in orders (Fig. 4) and families (Extended Data Fig. 6), metazoan classes behave differently than families and orders. Rather than cycling and turning over, the suite of classes we observe in the modern biosphere is dominated by the same classes that have existed since the Cambrian. Very few modern marine classes are “young.” In fact, of the major metazoan classes that are thoroughly represented in the PBDB, only 4 have ever gone extinct: Conodonta, Trilobita, Strophomenata, and Saurischia (which is non-marine). At the beginning and the end of the time series, splitting by “old” and “young” classes is therefore largely meaningless, as almost all of them belong to the same category. This problem is clearly displayed in Extended Data Fig. 8e-f.

**Extended Data Fig. 8:**
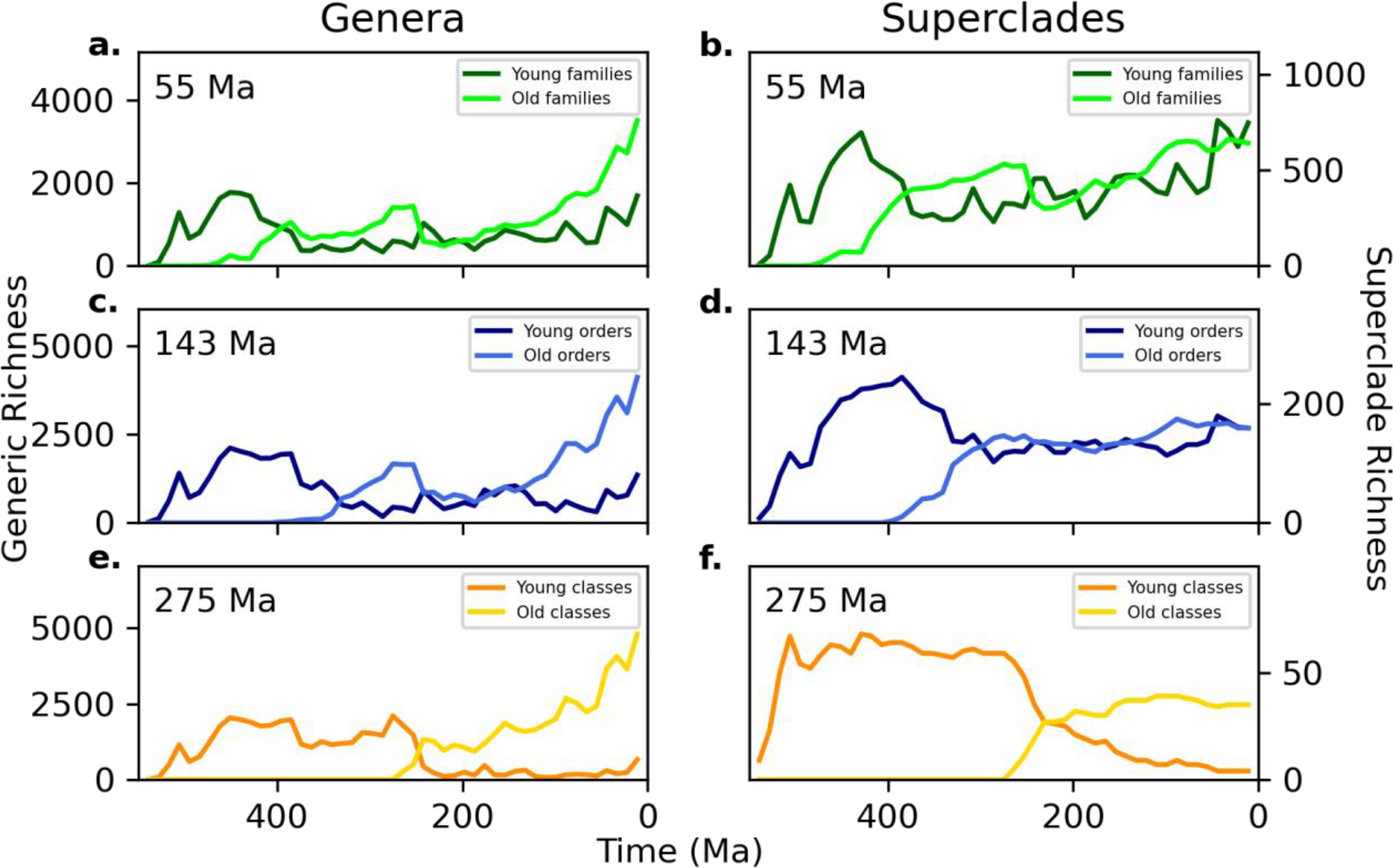
Number of genera in “old” and “young” superclades and the number of such superclades (a-b: families, c-d: orders, e-f: classes) plotted over time, as defined by the cut-offs in each top-left corner.

**Extended Data Fig. 9:**
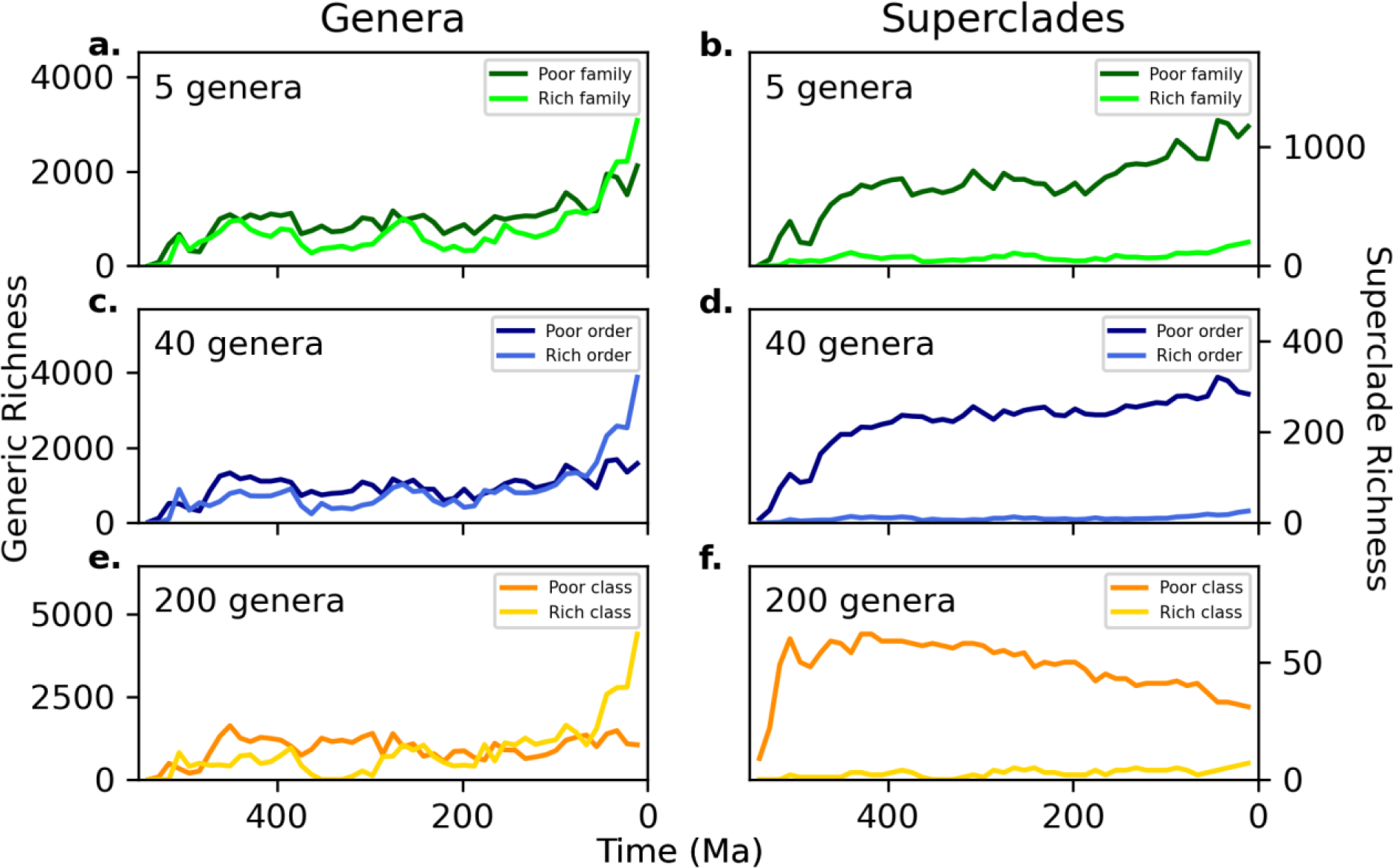
Number of genera in “rich” and “poor” superclades and the number of such superclades (a-b: families, c-d: orders, e-f: classes) plotted over time, as defined by the cut-offs in each top-left corner.

## References

[1] J. E. Lovelock, “Gaia as seen through the atmosphere,” Atmospheric Environment (1967), vol. 6, p. 579–580, 8 1972.

[2] W. F. Doolittle, “Is Nature really motherly?,” CoEvol. Quart. Spring, pp. 58–63, 1981.

[3] J. W. Kirchner, “The Gaia hypothesis: Can it be tested?,” Reviews of Geophysics, vol. 27, p. 223, 1989.

[4] J. W. Kirchner, “The Gaia Hypothesis: Fact, Theory, and Wishful Thinking,” Climatic Change, vol. 52, p. 391–408, 2002.

[5] W. F. Doolittle, “Natural selection through survival alone, and the possibility of Gaia,” Biology & Philosophy, vol. 29, p. 415–423, 6 2013.

[6] W. F. Doolittle, “Darwinizing Gaia,” Journal of Theoretical Biology, vol. 434, p. 11–19, 12 2017.

[7] W. F. Doolittle, “Making Evolutionary Sense of Gaia,” Trends in Ecology & Evolution, vol. 34, p. 889–894, 10 2019.

[8] T. M. Lenton, S. J. Daines, J. G. Dyke, A. E. Nicholson, D. M. Wilkinson and H. T. P. Williams, “Selection for Gaia across Multiple Scales,” Trends in Ecology & Evolution, vol. 33, p. 633–645, 8 2018.

[9] R. Arthur and A. Nicholson, “Selection principles for Gaia,” Journal of Theoretical Biology, vol. 533, p. 110940, 2022.

[10] A. E. Nicholson, D. M. Wilkinson, H. T. P. Williams and T. M. Lenton, “Alternative mechanisms for Gaia,” Journal of Theoretical Biology, vol. 457, p. 249–257, 11 2018.

[11] R. A. Boyle and T. M. Lenton, “The evolution of biogeochemical recycling by persistence-based selection,” Communications Earth & Environment, vol. 3, 3 2022.

[12] J. Alroy, C. R. Marshall, R. K. Bambach, K. Bezusko, M. Foote, F. T. Fürsich, T. A. Hansen, S. M. Holland, L. C. Ivany, D. Jablonski, D. K. Jacobs, D. C. Jones, M. A. Kosnik, S. Lidgard, S. Low, A. I. Miller, P. M. Novack-Gottshall, T. D. Olszewski, M. E. Patzkowsky, D. M. Raup, K. Roy, J. J. Sepkoski, M. G. Sommers, P. J. Wagner and A. Webber, “Effects of sampling standardization on estimates of Phanerozoic marine diversification,” Proceedings of the National Academy of Sciences, vol. 98, p. 6261–6266, 5 2001.

[13] S. Finnegan, J. L. Payne and S. C. Wang, “The Red Queen revisited: reevaluating the age selectivity of Phanerozoic marine genus extinctions,” Paleobiology, vol. 34, p. 318–341, 2008.

[14] M. Foote, “Survivorship analysis of Cambrian and Ordovician trilobites,” Paleobiology, vol. 14, p. 258–271, 1988.

[15] G. F. Boyajian, “Phanerozoic trends in background extinction: Consequence of an aging fauna,” Geology, vol. 14, pp. 955–958, 11 1986.

[16] G. E. Boyajian, “Taxon age and selectivity of extinction,” Paleobiology, vol. 17, p. 49–57, 1991.

[17] N. L. Gilinsky, “Survivorship in the Bivalvia: comparing living and extinct genera and families,” Paleobiology, vol. 14, p. 370–386, 1988.

[18] L. Van Valen, “A new evolutionary law,” Evolutionary Theory, vol. 1, pp. 1–30, 1973.

[19] T. C. Foin, J. W. Valentine and F. J. Ayala, “Extinction of taxa and Van Valen’s law,” Nature, vol. 257, p. 514–515, 1975.

[20] D. M. Raup, “Taxonomic Survivorship Curves and Van Valen’s Law,” Paleobiology, vol. 1, p. 82–96, 1975.

[21] J. J. Sepkoski, “Stratigraphic Biases in the Analysis of Taxonomic Survivorship,” Paleobiology, vol. 1, p. 343–355, 1975.

[22] S. N. Salthe, “Some Comments on Van Valen’s Law of Extinction,” Paleobiology, vol. 1, p. 356–358, 1975.

[23] A. R. McCune, “On the Fallacy of Constant Extinction Rates,” Evolution, vol. 36, p. 610, 5 1982.

[24] M. J. Benton, “The Red Queen and the Court Jester: Species Diversity and the Role of Biotic and Abiotic Factors Through Time,” Science, vol. 323, p. 728–732, 2 2009.

[25] A. Z. Krug, D. Jablonski and J. W. Valentine, “Species–genus ratios reflect a global history of diversification and range expansion in marine bivalves,” Proceedings of the Royal Society B: Biological Sciences, vol. 275, p. 1117–1123, 2 2008.

[26] M. E. Clapham and P. R. Renne, “Flood Basalts and Mass Extinctions,” Annual Review of Earth and Planetary Sciences, vol. 47, p. 275–303, 5 2019.

[27] N. J. Butterfield, “Animals and the invention of the Phanerozoic Earth system,” Trends in Ecology & Evolution, vol. 26, p. 81–87, 2 2011.

[28] Intergovernmental Panel on Climate Change (IPCC), Climate Change 2013 – The Physical Science Basis: Working Group I Contribution to the Fifth Assessment Report of the Intergovernmental Panel on Climate Change, Cambridge University Press, 2014.

[29] M. Schobben and B. van de Schootbrugge, “Increased Stability in Carbon Isotope Records Reflects Emerging Complexity of the Biosphere,” Frontiers in Earth Science, vol. 7, 5 2019.

[30] A. Calbet, “Mesozooplankton grazing effect on primary production: A global comparative analysis in marine ecosystems,” Limnology and Oceanography, vol. 46, p. 1824–1830, 11 2001.

[31] A. Chopra and C. H. Lineweaver, “The Case for a Gaian Bottleneck: The Biology of Habitability,” Astrobiology, vol. 16, p. 7–22, 1 2016.

[32] F. Pedregosa, G. Varoquaux, A. Gramfort, V. Michel, B. Thirion, O. Grisel, M. Blondel, P. Prettenhofer, R. Weiss, V. Dubourg, J. Vanderplas, A. Passos, D. Cournapeau, M. Brucher, M. Perrot and E. Duchesnay, “Scikit-learn: Machine Learning in Python,” Journal of Machine Learning Research, 10 2011.

